# Cell type-specific interpretation of noncoding variants using deep learning-based methods

**DOI:** 10.1101/2021.12.31.474623

**Authors:** Maria Sindeeva, Nikolay Chekanov, Manvel Avetisian, Nikita Baranov, Elian Malkin, Alexander Lapin, Olga Kardymon, Veniamin Fishman

## Abstract

Interpretation of non-coding genomic variants is one of the most important challenges in human genetics. Machine learning methods have emerged recently as a powerful tool to solve this problem. State-of-the-art approaches allow prediction of transcriptional and epigenetic effects caused by non-coding mutations. However, these approaches require specific experimental data for training and can not generalize across cell types where required features were not experimentally measured. We show here that available epigenetic characteristics of human cell types are extremely sparse, limiting those approaches that rely on specific epigenetic input. We propose a new neural network architecture, *DeepCT*, which can learn complex interconnections of epigenetic features and infer unmeasured data from any available input. Furthermore, we show that DeepCT can learn cell type-specific properties, build biologically meaningful vector representations of cell types and utilize these representations to generate cell type-specific predictions of the effects of non-coding variations in the human genome.

---

During the development of a multicellular organism, a single cell gives rise to a large diversity of cell types. These cell types dramatically differ from each other, although sharing almost identical DNA sequences. For example, the same promoter sequence may drive drastically different expression levels of a downstream gene depending on cell type. These differences in expression stem from two sources. First, sequence-specific binding by trans-factors, which are expressed in one cell type but not another, controls transcriptional activity. Second, some loci became epigenetically bookmarked at the stage of progenitor cells, making these loci unresponsive to trans-factors expressed later in development. See (Heinz et al. 2015) for detailed review.

Developmental history and the presence of trans-factor together define the *cell state*, explaining differences in epigenetic properties and transcription levels of the same sequence. Cells sharing the same or close states represent a cell type. Due to differences in cell state, genomic variations may have different, cell state-specific transcriptional or epigenetic effects. This makes interpretation of genomic variations, including clinical assessments of non-coding mutations, challenging.

Importantly, cell states could be learned from the genome-wide expression pattern (Nair et al. 2019; Cinghu et al. 2014) or epigenetic marks (Schreiber et al. 2020) allowing clustering of similar cell types based on their genomic properties. Moreover, many epigenetic marks are interdependent (Ernst and Kellis 2015), making it possible to use only a subset of measured epigenetic characteristics to infer unmeasured properties and define cell state. Due to the complex nature of dependencies between epigenetic marks machine learning approaches recently gained attention as attractive tools to solve this problem (Wong et al. 2021); however, the existing solutions (Keilwagen, Posch, and Grau 2019) focus on a specific combination of input and target epigenetic properties, lacking generalizability to infer all epigenetic properties from any available input.

Although the cell state is essential to explain differences between cell types, it could not explain differences in the expression levels or chromatin packaging associated with different sequences within the same cell type. To explain these differences, one should consider the diversity of DNA sequences. There are several computational methods allowing the prediction of specific epigenetic properties (Chen et al. 2021; Avsec et al. 2021), chromatin packaging (P. Belokopytova and Fishman 2021), or expression levels (Avsec et al. 2021) based on the DNA sequences. Among these, the prediction of gene expression changes caused by sequence variations is especially interesting, because such predictions enable clinical interpretation of non-coding variants in the human genome (Avsec et al. 2021). However, none of the existing predictive approaches allow generalization of the predictions across cell types, limiting the results to a subset of cell types with known expression patterns.

Here, we introduce the concept of cell state learning, which allows us to build latent vector representations of cell types based on available epigenetic input. We show that sequence specificity could be combined with cell type-specificity using modern machine learning architectures. We show the feasibility of the developed models for the cell type-specific prediction of epigenetic properties. Importantly, we showed that these predictions allow inferring functional significance of effects caused by noncoding variants in the human genome.

## Results

### AI model allows learning meaningful computer representation of cell types using arbitrary epigenetic data

We started our investigation by exploring the latest update of the ENCODE dataset (Boix et al. 2021). Focusing on human data, we collected information about 858 cell types characterized by 40 epigenetic features, 3026 tracks in total (Fig. 1). Among these epigenetic features, measurements of DNAse accessibility are the most abundant (available for 631 cell types), and the less-studied marks are H2AK9ac, H4K12ac, and H3T11ph, which are measured only in one cell type (Fig. 1, A). The most characterized cell type is IMR-90 fibroblasts, with 33 out of 40 epigenetic tracks available; at the same time, for 464 cell types, only one epigenetic mark was measured (Fig. 1, B). Overall, we found that available information about epigenetic characteristics of human cells is sparse, with only 8.8% of epigenetic tracks present in the ENCODE collection.

**Figure 1.**
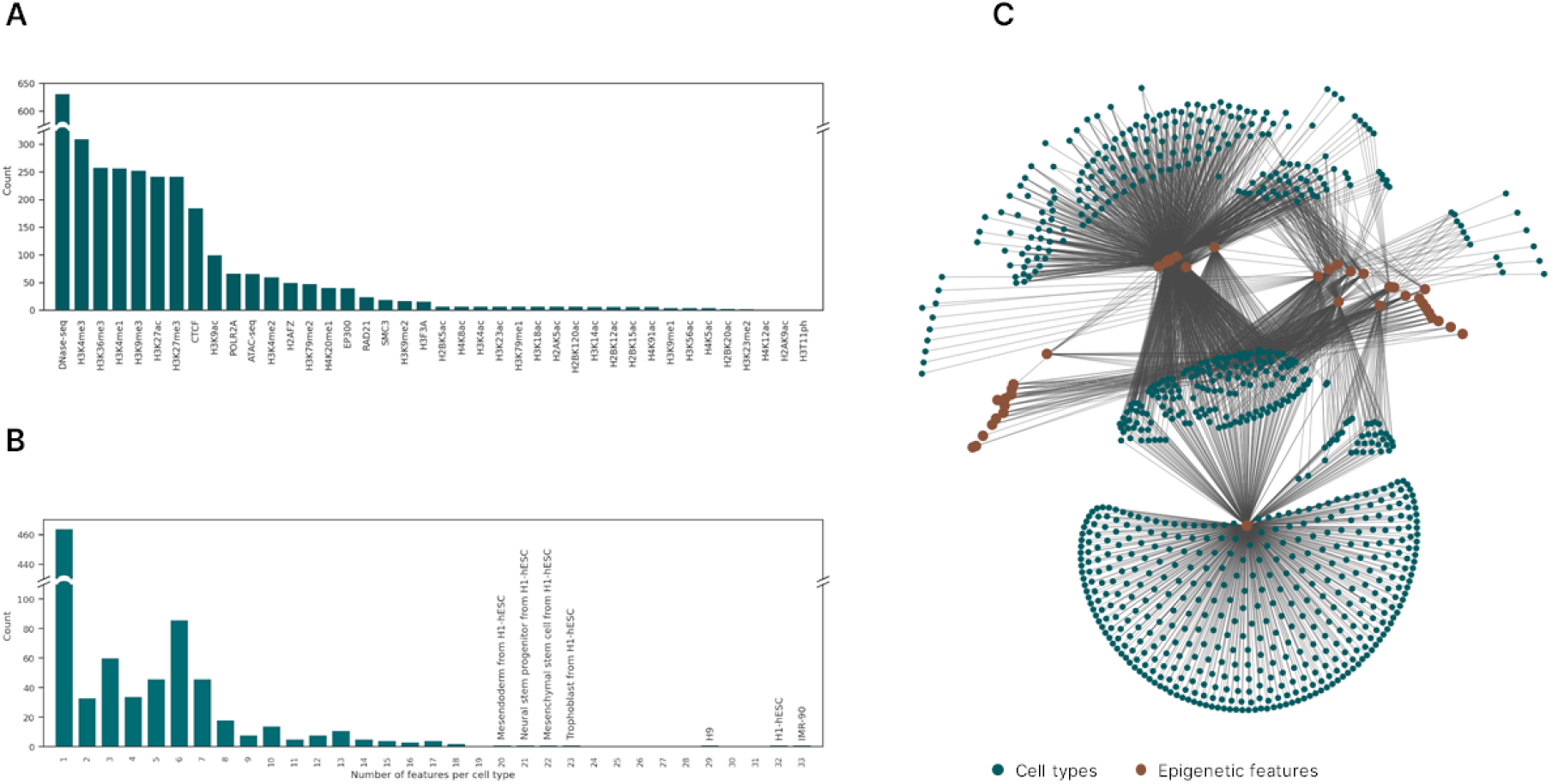
The overview ENCODE epigenetic datasets characterizing human cell types. A - number of cell types specific epigenetic profiles shown for each epigenetic feature. B - histogram showing a number of epigenetic features measured per cell type. C. Graph representation of measured epigenetic data. Each node presents cell type (green nodes) or epigenetic features (brown nodes), each edge shows measured cell type-feature track.

We made an assumption that if two epigenetic features were measured in one cell type, it is possible to learn how the signal of one feature depends on the signal of another feature. We also assumed that dependencies between epigenetic features are largely shared between cell types. This assumption might not hold if there are cell type-specific epigenetic mechanisms. However, measured data should allow clustering of epigenetically similar cell types, which probably share mechanisms of epigenetic regulation. Within these clusters, we can utilize dependence between signals of two features learned in one cell type to predict the signal of an unmeasured feature in another cell type.

To explore how often unmeasured features can be inferred from measured data, we visualized and analyzed cell types and features as a graph where edges represent feature measurements (Fig. 1, C). We found that this graph is connected, therefore under the aforementioned assumptions, any unmeasured feature can be inferred from the available data. Thus, we aimed to develop a computational approach capable of clustering epigenetically similar cell types and infer unmeasured epigenetic features based on available sparse datasets. Towards this aim, we decided to employ state-of-the-art machine learning methods.

We have designed new neural network architecture, *DeepCT*, capturing both sequence and cell type-specific variations of the epigenetic data (Fig 2, A). The model accepts DNA sequence and cell type label as its inputs. The DNA tail processes the nucleotide sequence and builds its computational representation using a deep convolutional neural network (CNN). The cell state tail accepts a one-hot encoded vector of the input cell type and returns the embedding vector which represents this cell type in the model’s latent space. The sequence and cell state representations are concatenated and fed to the network’s head, which outputs sequence- and cell state-specific predictions about the presence of each epigenetic mark.

**Figure 2.**
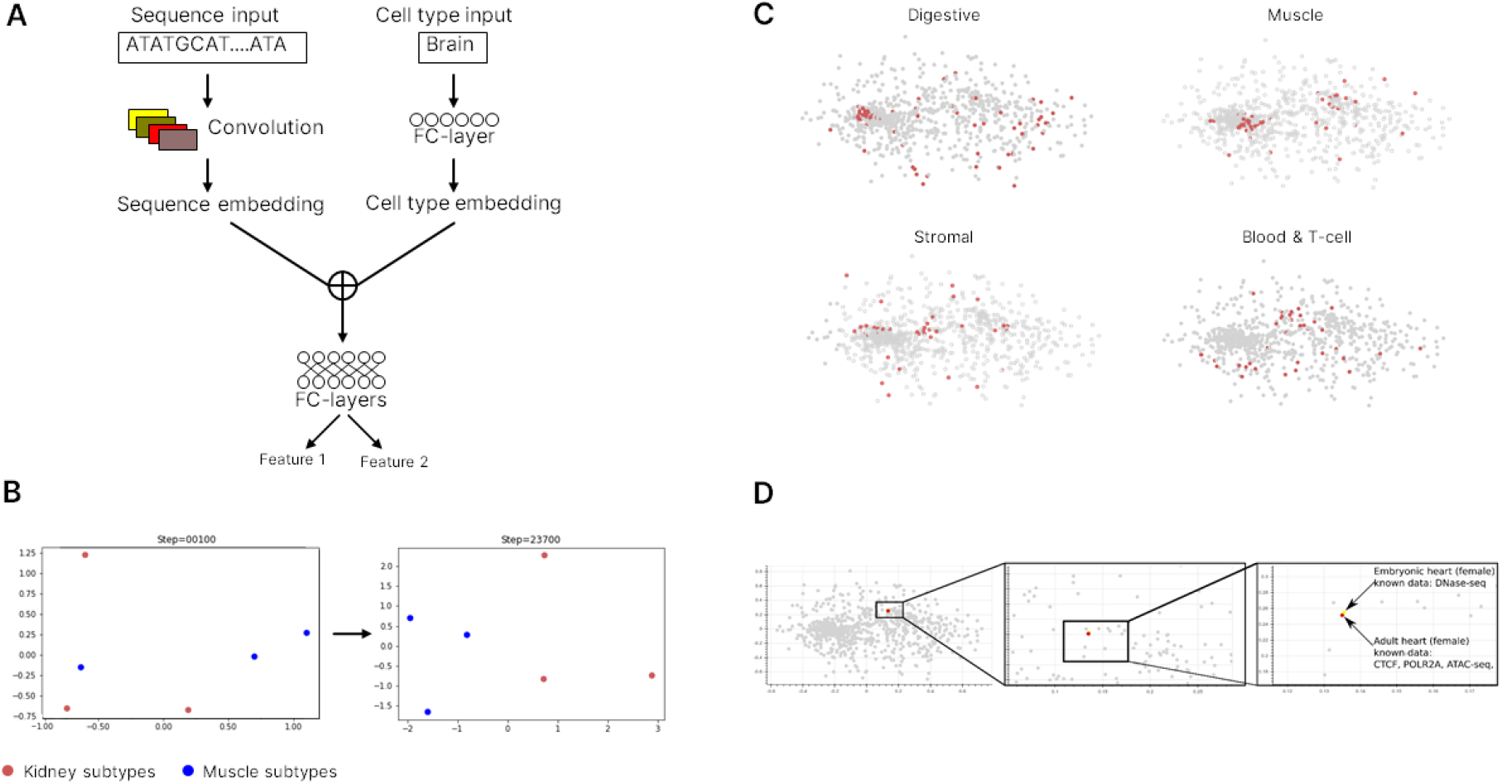
DeepCT models learn computer representations of cell type. A. An overview of the DeepCT models architecture. Details are provided in methods and Supplementary Note 1. B. Projection of embeddings for three kidney and three muscle cell types from 32-dimensional latent space of the model into 2-dimension PCA axis. The figure shows how cell states, initialized randomly, are clustered according to cell identity after ∼23 000 model training steps. C. Example cell state embeddings projections from 32-dimensional latent space of the model into dimension PCA axis. Background points show distribution of embeddings for all cell types, whereas colored dots correspond to the specific cell types associated with specific tissue: digestive, muscle, stromal or blood. D. DeepCT performs clustering of embryonic and adult heart tissues, although there is no common epigenetic track measured for both samples,

The idea of cell state learning described above could be implemented using different neural network architectures. We demonstrated this by designing several implementations of the DeepCT model (methods). The qualitative implementation (DeepCT) aims to solve classification problems, i.e. the presence or absence of a peak at a specific locus in a specific cell type. The quantitative representation (qDeepCT) solves the regression problems, predicting the ChIP-seq, ATAC-seq, DNAse I hypersensitivity or CAGE data quantitatively. The quantitative implementations use CNN for sequence processing (CNN-qDeepCT and trans-qDeepCT). Following, we provide results for CNN-qDeepCT model, because it shows better performance on our data.

We expected that during training, the network will fit its parameters so that similar sequences obtain similar embeddings and, likewise, similar cell types will be clustered within the model’s latent space. Indeed, a toy example of six cell types characterized by a single epigenetic feature shows how clustering of similar (sub)types occurs during neural network training (Fig. 2, B; Supplementary Video S1).

Applying DeepCT to the full dataset (excluding data obtained after treating cells with specific compounds) of 2629 high-quality tracks, we observed that cell type clustering was often concordant with biological expectations (Fig. 2, C, D; Supplementary Fig. 1). For example, we show co-localization of muscle cells, as well as co-localization of digestive cells (Fig. 2 C). However, other cell types, such as blood or stromal cells do not form well-defined clusters (Fig. 2, C). This might be either due to higher heterogeneity of these tissues or due to the peculiar properties of the epigenetic data available for these cell types.

To quantify the performance of clustering, we measured how far the cell types of the same tissue or organ appeared to be located within the model’s latent space (Supplementary Fig. 1). This analysis confirmed that average cosine similarity for embeddings representing cell types from the same tissue was significantly higher than for embeddings of randomly selected cell types (0.229 vs 0.176 ± 0.004 at random). This result shows that the DeepCT model captures the biological properties of cells during training.

We note that DeepCT learned the cell representation based on different collections of epigenetic marks available for each cell type. For example, we observed clustering of embryonic and adult heart samples (Fig. 2, D), although the set of epigenetic tracks measured in adult tissues (CTCF and POLR2A ChiP-seq and ATAC-seq) does not contain a single track available for embryonic heart (DNAse I hypersensitivity). This is possible because all available epigenetic data are fed to one model, sharing the latent space, and thus allowing multitask training. In this example, we may speculate that ATAC-seq and DNAse I hypersensitivity are highly correlative, which was learned from those cell types where both features are available. This correlation can be utilized to infer ATAC-seq data in embryonic heart and cluster embryonic and adult tissues.

Thus, using the developed architecture allows the cell type embeddings to be fitted using any subset of the epigenetic datasets. These embeddings can be next used to infer unmeasured features.

### DeepCT model generalizes across cell types, sequences, and epigenetic tracks

To benchmark the developed DeepCT models, we considered three challenges (Fig 3, A-C). For the first, cell type specificity challenge, we excluded 10% of epigenetic data for each locus and evaluated the model on these excluded targets (Fig 3, A; see also details in methods). Note that in this setup for each locus the training data includes targets measured for some, but not all, cell types. As a baseline in this challenge, we used the average target value for a given locus across cell types included in training. Biologically, this baseline can be interpreted as an average cell type’s epigenetic profile. We note that this baseline has performance metrics substantially higher than expected in random (baseline AP=0.417), because differences between sequences explain substantially more variance in epigenetic data than differences between cell types.

**Figure 3.**
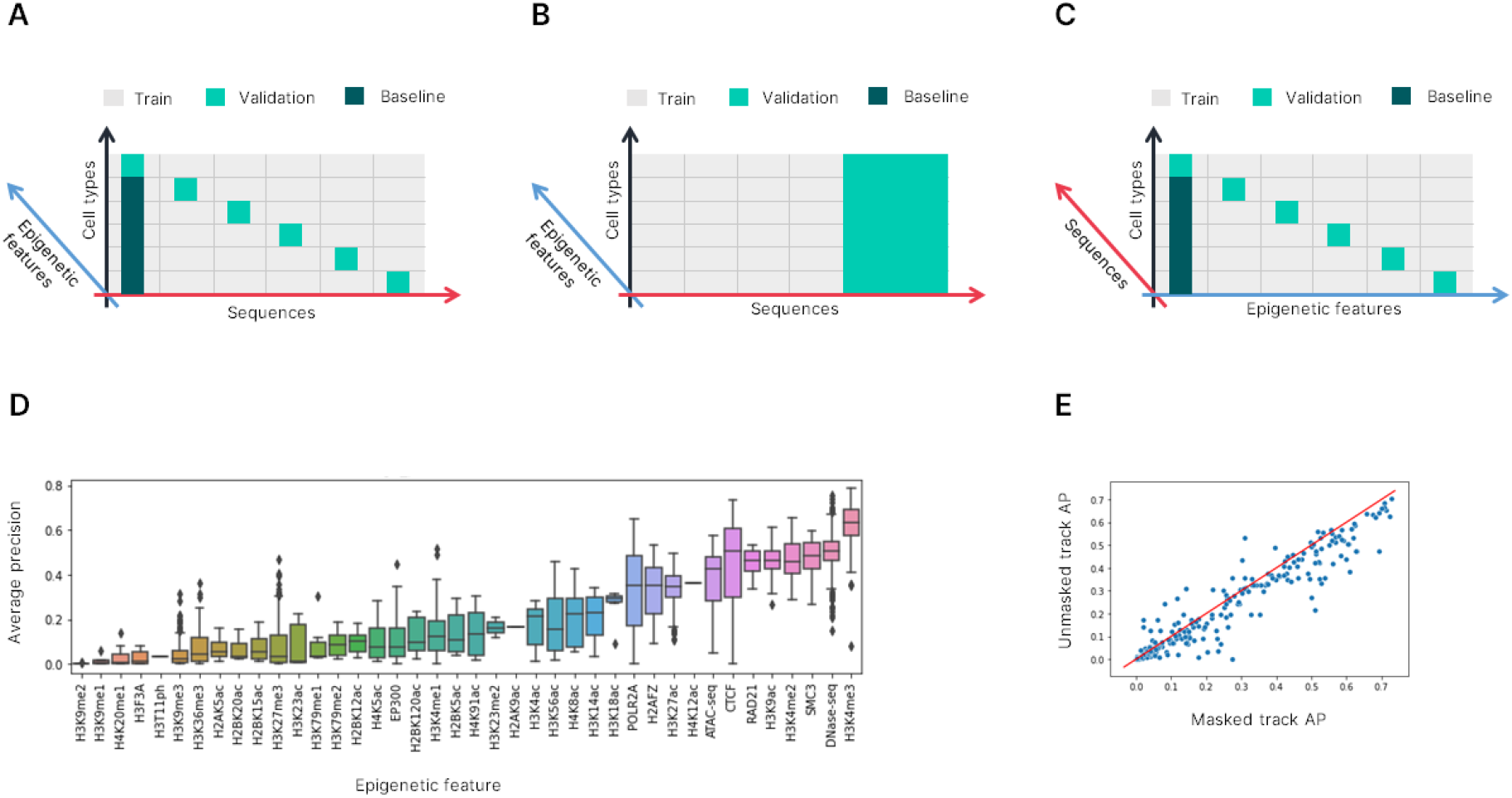
Benchmarking DeepCT architecture. A-C. Different schemes of DeepCT benchmarks. We show three dimensions of the target epigenetic signal values predicted by the neural network: sequence dimension (representing diversity of genomic loci), cell type dimension and epigenetic feature dimension. Each target value is defined by a combination of genomic loci, epigenetic feature and cell type. Colors show which targets were excluded from training data. Note the axis swap in panel C: in A and B we use at least 50% of examples for any cell type x feature combinations for model training, whereas for C we completely removed data for some of the cell type x feature combinations from the training dataset. D. Distributions of average precision (AP) values obtained for different cell types. AP measured for unseen sequences. E. Scatter plot showing how AP changes when excluding epigenetic track from training. X-axis shows AP measured for a track included in training (i.e. track’s data was included for training sequences set). Y-axis shows AP measured for the same track, but in this case, no track’s data was included during training, requiring the model to infer the track values from the cell state embedding. In both cases, AP measured for the same set of unseen sequences.

We compared performance of the DeepCT models with baseline metrics and found that the model outperforms the baseline (model AP 0.430 vs baseline AP 0.417), showing that cell state embeddings are meaningful and that the model predicts cell type-specific differences in epigenetic features using learned embeddings.

Next, we considered the sequence specificity benchmark. In this test, we defined a subset of validation loci and completely excluded any information about these loci from the training dataset. This is a more complicated challenge compared to the previous one when for each sequence, a subset of target values was available during training. Expectedly, the model’s performance was lower in this case (mean AP ∼0.34, mean r2 ∼0.16). However, the performance significantly varies among features (Fig. 3, D). For example, H3K9 and H4K20 monomethylation have almost zero average precision; on the other hand, H3K4me3 mark shows high AP score (mean across cell types 0.62), and for best cell types we achieved AP ∼0.8 on unseen sequences. The predictive power was also high for DNAaseI hypersensitivity, CTCF, Cohesin subunits, and some other epigenetic datasets.

Finally, the most complicated challenge is the prediction of previously unmeasured epigenetic tracks. To benchmark our model, we excluded a subset of tracks from the training dataset. We selected these tracks so that each cell type was characterized by at least one epigenetic track, allowing us to learn an embedding for this cell type. In addition, we ensured that each excluded track can be inferred by learning dependencies between epigenetic features from the training data (methods).

Expectedly, the quality of the predictions was lower for unseen tracks (Fig. 3, E). However, the drop of predictive power was relatively small, only 1.6 % of AP score on average. This indicates that DeepCT architecture can be used to infer previously unmeasured data, serving as an important instrument for interpretation of non-coding variants in the human genome.

#### Using the DeepCT models for interpretation of clinically significant genomic variants

To benchmark the clinical relevance of the obtained predictions we employed the collection of genomic variants previously associated with autism spectrum disorder (An et al. 2018; Zhou et al. 2019). Although epigenetic impact was recently measured for a large collection of non-coding variants (Abramov et al. 2021), for the vast majority of variants identified by (An et al. 2018) and (Zhou et al. 2019) epigenetic effects were not known. To gain this information, we predicted 40 epigenetic characteristics for each single nucleotide variant (SNV) in the dataset. Comparing predictions obtained for reference and alternative allele, we scored the effects of SNVs identified in children with autism spectrum disorder and their unaffected siblings.

We found that effects of variants identified in genomes of children with autism spectrum disorder were on average larger than effects of variants carried by their unaffected siblings (Fig. 4, A, B). This difference was observed both for epigenetic tracks where reference alleles were experimentally characterized and for epigenetic tracks that were never measured before. We selected genomic variants showing the largest effects (top 500 of SNVs or ∼1% of all SNVs analyzed) and assigned the nearest gene to each SNV. Ontology analyses of the obtained genes set showed significant enrichment of regulation of GTPase activity, neuron projection development, neurogenesis, and other relevant processes (Supplementary Table 1).

**Figure 4.**
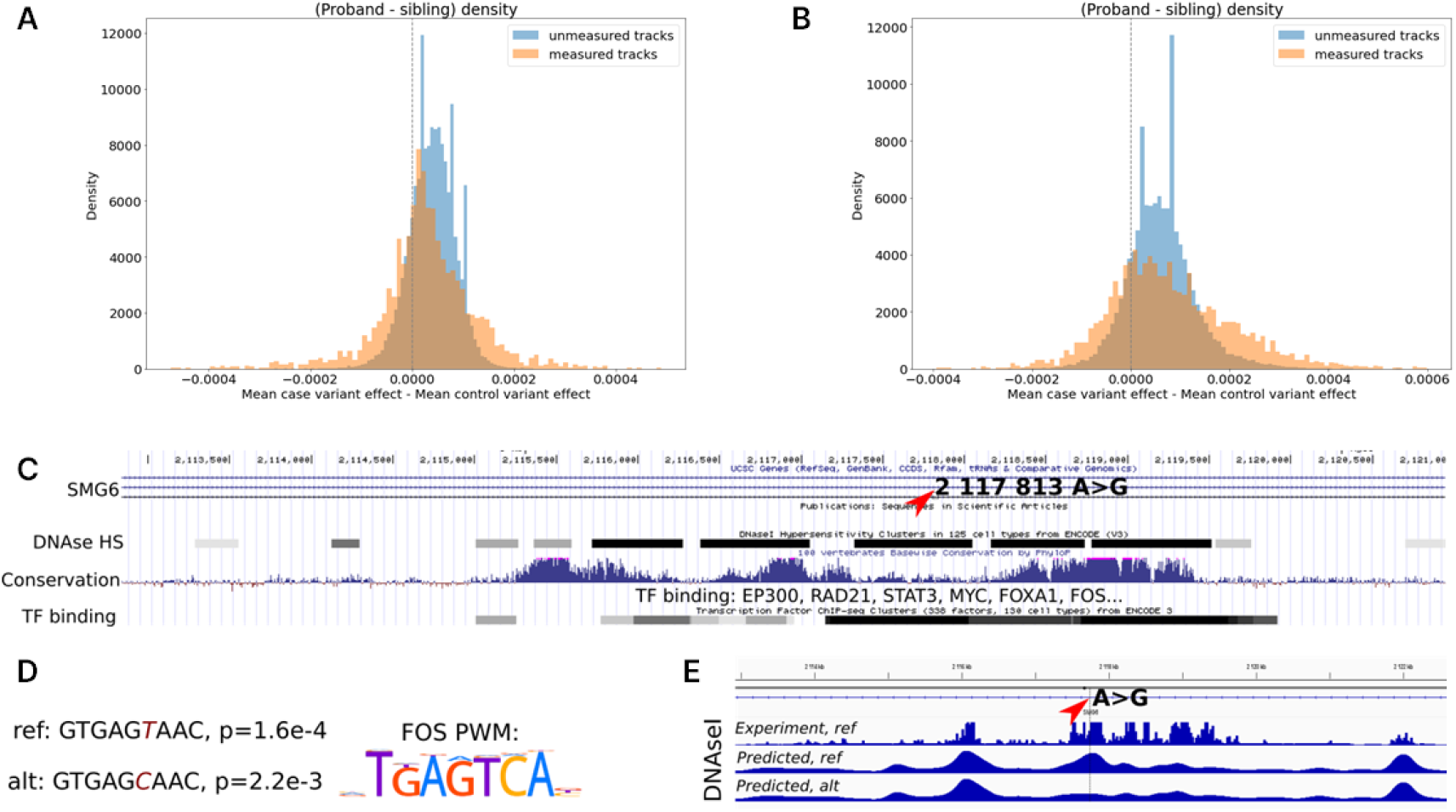
DeepCT predicts effects of SNVs associated with autism spectrum disorders. A and B. Distribution of mean difference of effect size for affected child’s and unaffected sibling SNVs. Data is average across SNVs, i.e. each sample in the distribution corresponds to one of 31 760 epigenetic tracks (794 cell types x 40 epigenetic features) predicted by DeepCT. Datasets where experimental measurement for reference allele was available during model’s training is highlighted as “measured tracks”. Effects of SNVs from (An et al. 2018) shown in A, for (Zhou et al. 2019) shown in B. C. Genomic regions within SMG6 gene intron containing NC_000017.10:g.2117813A>G variant (depicted by arrow). D. FOS motif p-values for reference and alternative alleles computed based on FOS position weights matrix (PWM). Note that we show reverse complement of the sequence matching PWM. E. DNAseI hypersensitivity tracks for M059J cell line. Position of NC_000017.10:g.2117813A>G variant depicted by arrow.

We focused on specific SNVs with high effect sizes. These SNVs often fall into DNAse I hypersensitive regions with multiple binding sites for key developmental regulators. For example, we identified an SNV within the intron of *SMG6* gene (NC_000017.10:g.2117813A>G), approximately 50 Kbp from its transcription start site (Fig. 4, C). Previously, single nucleotide variants and copy-number variations affecting these genes were associated with autism spectrum disorders (Shi et al. 2013; Nguyen et al. 2013; Iossifov et al. 2014). Consistently, DeepCT predictions show the highest impact for the M059J line, which originates from glial cells.

Although there were other SNVs identified within the *SMG6* gene, only this SNV was scored high by DeepCT, predicting almost ten-fold change of DNAse I accessibility. In accord with high impact predicted by DeepCT, this SNV falls within an evolutionary conserved genomic locus, demarcated by H3K27 acetylation marks and containing multiple transcription factor binding sites (Fig. 4, C). Analysis of transcription factor motifs based on HOCOMOCO motif search tool (Kulakovskiy et al. 2018) shows that the SNV significantly decreases the probability of FOS transcription factor grinding in this region (Fig. 4, D). Accordingly, DeepCT predicts substantial decrease of DNAse I sensitivity for alternative allele (Fig. 4, E) These observations can explain how this SNV leads to dysregulation of *SMG6* gene expression and contribute to the development of the autism spectrum disorder.

## Discussion

We challenged one of the most important questions in modern genomics: interpretation of non-coding genomic variants. We show how this question can be answered using state-of-the-art machine learning methods. For this aim, we decomposed epigenetic profiles into two components: one representing cell type specific mechanisms (cell states) and another representing diversity of genomic sequences. Speaking in biological terms, we can imagine that two heads of DeepCT network are responsible for learning mechanisms of *cis-* and a *trans-* regulation, whereas the tail of DeepCT learns interplay between these two factors and their epigenetic manifestations.

Although the possibilities opened by DeepCT-based neural network architectures seem promising, we would like to discuss limitations of our approach and directions toward improvement of DeepCT-based architectures.

First, we can not guarantee generalizability of the model across all human cell types. In fact, we expect that very specific epigenetic profiles, observed, for example, in germ cells, zygote (Du, Zhang, and Xie 2021), or some adult cell types, such as erythrocyte progenitors (Ryzhkova et al. 2021; Fishman et al. 2019) will not be inferred by DeepCT. In addition, in this work, we did not control how the cell types were split between training and validation sets. This may lead to inflated performance when two very similar cellular subtypes appear in train and validation datasets. On the other hand, there is currently no “ground true” broad classification of ENCODE cell types which would reflect their epigenetic similarity. The best classification we were able to find (presented in Supplementary Fig. 1) is derived using ENCODE cell ontology database, reflecting anatomical rather than epigenetic relationships between different cell types. In fact, we hope that the latent space of cell states inferred by DeepCT can be used to build the best epigenetic classification of human cells in the future, although there are some limitations of this approach discussed below.

Although we use the term *cell type* through the manuscript, in this work we employed multiple epigenetic profiles of heterogeneous tissues, composed of different cell types. Applying DeepCT to more pure single-cell datasets can extend cell types classification and may help to solve an important challenge of learning missing modalities from single cell data.

Although we show that cell type representations learned by DeepCT model are meaningful, we emphasize that these representations include a component associated with technical biases of experiments. Indeed, cell state embedding is learned to explain all dependencies between measured epigenetic profile and sequence composition specific for particular cell type. This specificity may originate not only from biological ground, but also from poor quality of experimental conditions specific for this cell type. Of course, with a large number of experimental datasets available for one cell type (ideally obtained in different laboratories to reduce batch-effect) this factor would play a lesser role in resulting embeddings.

Finally, we note that performance of the DeepCT model for some tracks and some sequences was low, although in other cases we obtained high performance scores. We expect that performance of DeepCT can be further improved using more powerful neural network architectures (such as transformer layers, (Avsec et al. 2021)), including information about long-range regulation (P. Belokopytova and Fishman 2021), and including more epigenetic datasets.

## Methods

### Data collection

#### ENCODE data collection

We downloaded re-processed ENCODE datasets from EpiMap (Boix et al. 2021) *http://compbio.mit.edu/epimap/*. We used p-value rather than fold-change as a signal measure, motivated by suggestions from (Boix et al. 2021). The complete datasets contained 3026 tracks, i.e. cell type / epigenetic feature combinations. For model training, we excluded tracks with low-quality tracks (the list was downloaded from supplementary information available in (Boix et al. 2021). The resulting dataset includes 2629 tracks describing 40 epigenetic features (41 if CAGE data included; see below) characterizing 794 human cell types.

To convert the quantitative p-value track into qualitative peaks positions, we used an empirical threshold of negative log of p-value equal to 4.4 (i.e. p-value below 10^−4.4^). We defined peaks as 150 bp genomic intervals centered around genomic positions where the signal exceeds the threshold. The threshold value and peak length were chosen to match peak positions reported by standard Encode processing pipelines. Overlapping peaks were merged, and only those intervals where at least one peak was detected were kept in the dataset. For training the quantitative model, we logged data and clipped values using thresholds -1.0…4.0.

#### CAGE dataset collection and processing

We used metadata from the GTRD (Kolmykov et al. 2021) to match ENCODE with FANTOM5 (Forrest et al. 2014). Primarily we used *cell_id* identifier from the GTRD which uniquely describes cell type; in addition, we manually screened results to ensure the same origin and treatment status, removed fetal samples (as expression and epigenetic status may rapidly change during the development and there’s no way to ensure a proper match between samples from different organisms even at exact same-day age—which by itself almost never matches between these databases), and corrected annotation mistakes. We downloaded alignments for chosen experiments from *https://fantom.gsc.riken.jp/5/datafiles/latest/basic/*; data from biological replicates were merged together. Lastly, we connected FANTOM5 experiments to EpiMap’s *BSSID* identifiers which describe cell type, age and treatment status; in case of 1-to-many and many-to-many relationships we used random assignment to create pairs. The resulting dataset comprised 99 pairings.

To obtain coverage track we processed alignments with DeepTools bamCoverage using RPKM normalization, bin size of 10, smoothing window size of 50, and minimum mapping quality threshold of 1. For training, we logged data and clipped values using thresholds -7.0…2.0. Note that we used CAGE data only for quantitative models.

### Machine learning models architecture

#### Models inputs and targets

We constructed CNN models with two inputs: a sequence input, accepting one-hot encoded 1000 bp DNA sequence (shape 4×1000), and a cell type input, accepting one-hot encoded cell type (shape 1×N_cell_types). We defined a 200 bp region in the center of the input sequence as a target region, and constructed the model’s target as follows:

a. For qualitative models, we used binary labels 1 or 0; label 1 indicates that in the specified cell type and epigenetic feature combination, there is the epigenetic feature peak overlapping the selected 200 bp region by at least 50%.
b. For quantitative models, we used the maximal feature signal value computed across the target 200 bp region for all data except CAGE. For CAGE data, we used the sum of signals.

To sample sequences, we filtered out genomic intervals where no peak of target features is present. We extended the remaining intervals by 30 base pairs from both sides (to add samples with no peaks), and drew coordinates of centers of samples (sampling points) using uniform distribution. The sampling points were set apart by at least 200 base pairs, so that their targets would not overlap one another.

For a specific sequence input, we perform a forward pass of the model for all cell types at once to reduce computation load (see below). Thus, the model returns *N_cell_types* × *N_features* values, each value corresponding to a combination of cell type and epigenetic feature. It is important to note that even if some cell type / epigenetic feature combinations were not experimentally measured, the model outputs the prediction for these combinations, although these values did not contribute to the model’s loss (see below).

#### Models architecture

The models process DNA sequence by convolution tail, which outputs sequence embedding of shape 1×256. The layers specification could be found in Supplementary Note 1.

The cell state tail maps one-hot encoded cell type from the dimension R^N_cell_types^ to the dimension of cell type embeddings R^emb_length^ using a linear transformation (Supplementary Note 1). We set *emb_length* to 32.

To predict cell type-specific outputs, we concatenated sequence and cell type embeddings and passed the result to the model’s tail, which consists of eight fully connected layers (Supplementary Note 1). We set the number of the last layer’s outputs equal to the number of epigenetic features, thus for each sequence and cell type inputs we obtained predictions of all epigenetic features.

To reduce computational load, for each sequence we computed sequence embedding only once and then passed it to the model’s head concatenated with different cell type embeddings. Thus, for each sequence, we obtained *N_cell_types* × *N_features* predictions.

After the first trials, we found that predicting data for unseen sequences is substantially more challenging than for unseen cell types (see results for detail). These results fit with our observation that the portion of the variance in epigenetic data explained by sequences diversity is several times higher than the portion explained by cell types diversity. Motivated by these data, we decided to split the model’s head to provide two outputs: one providing sequence-specific prediction and another responsible for the cell-type-specific part of the prediction.

Let us consider a dataset of *N* cell types characterized by *K* epigenetic tracks. For the genomic position *s*, we aim to predict a set of target values *t*_*n,k,s*_, where *n* is a cell type index (*n=1*..*N*) and *k* is an epigenetic feature index (*k=1*..*K*). We defined sequence-specific mean feature positional information (MPI) as *M*_*k,s*_:

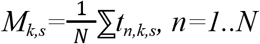

and cell type-specific deviations as *D*_*n,k,s*_:

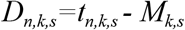

Thus, *M*_*k,s*_ is the mean feature value, which is invariant across cell types and depends on sequence only; *D*_*n,k,s*_ is cell type-specific deviation, it depends on cell type and shows how the particular cell type differs from average.

We used eight fully connected layers with final output shape (1×N_features) to predict *M* from sequence embedding. Similarly, we used eight fully connected layers with final output shape *N_cell_types* × *N_features* to predict *D* from concatenated sequence and cell type embeddings. Knowing *M* and *D*, we were able to reconstruct original targets simply by adding up *t*_*n,k,s*_ *= D*_*n,k,s*_ *+ M*_*k,s*_.

### Computing loss function

We used binary cross-entropy loss for qualitative models and MSE loss for quantitative models. For a model that outputs *M* and *D* values, we measured MSE loss as:

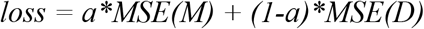

We found that *a* value equal to 0.0002 results in optimal performance. Interestingly, the main component of loss, in this case, is cell type-specific. Using a ∼ 0.5 significantly lowered performance metrics.

The model predicts target values for all cell type / feature combinations; however, for many combinations, there was no experimentally measured data, and therefore loss function can not be computed for outputs representing these combinations. Thus, only measured targets contribute to the loss. Similarly, we used only a subset of cell type/feature combinations that have experimentally measured data to compute *M* and *D* values.

### Training models

We trained models for 20 epochs using Tesla V100 16 GB GPU, with an initial learning rate set to 0.0001 and updated during training using a cosine annealing schedule, which we found to give the best model performance. We used selene-based framework for model training (Chen et al. 2019).

### Evaluation of models

#### Train and validation split

##### Unseen sequence prediction benchmark

For the unseen sequence prediction benchmark, we split our data by chromosomes, which prevents data leaks between overlapping regions (P. S. Belokopytova et al. 2020). We used two chromosomes for validation, two chromosomes for evaluation, and the rest of the genome to train models. Chromosome Y was excluded from all analyses.

##### Unseen cell-type prediction benchmark

For an unseen cell-type prediction benchmark, we wanted to see how well our model can differentiate between cell types. We did so by evaluating how well it can predict values for unseen combinations of cell types and sequences. We left out two chromosomes for model evaluation and split the rest of the chromosomes into several (3, 5, or 10) non-overlapping folds. We also split a set of cell types into a corresponding number of non-overlapping sets. We then chose to leave some (3, 5, or 10 respectively) pairs of these sequence and cell type subsets out of training for validation; the rest were kept for training, as demonstrated in Figure 3, A. Note that it would not be enough to simply leave some cell types out of training for validation, as the model would have no data to learn anything about these cell types. However, if we leave out pairs of (cell type, sequence), each cell type is still present in training, and thus we can evaluate whether the model’s predictions are cell-specific or only sequence-specific within sequence folds.

To ensure that all cell type folds contain the same number of tracks we chose the largest clique from the graph of measured tracks shown in Figure 1, C, and trained our models on a subset of tracks given by this clique. This left us with 905 tracks (181 cell types and 5 features).

To select tracks for unseen tracks prediction benchmark we performed the following steps. First, we filtered out all data for cell types represented by less than three features and for all features with less than three cell types experimentally profiled. Next, we removed all cell types with treatment. Finally, we aimed to remove 10% of tracks from the training dataset so that these tracks’ data could be predicted using the remaining tracks. For this aim, we defined track’s predictability: measurement of feature *X* in cell type *I* is predictable if and only if there is a cell feature *Y* and cell type *II* satisfying the conditions:

1. feature *Y* is measured in cell types *I* and *II*
2. feature *X* is measured in cell type *II*

We next randomly selected 10% of the tracks so that each of them could be predicted (based on the definition above) using the remaining tracks.

#### Performance metrics

To evaluate the qualitative model, we used the average precision (AP) metric. We pick this metric because it is robust for imbalanced data with numerous negative samples, which is the case for epigenetic datasets (see for example comparison presented in (Li, Quang, and Guan 2019)). To evaluate the quantitative model, we used the r^2^ metric. To compare results obtained by quantitative and qualitative models, we obtained peak positions from predicted quantitative signal using the same threshold as for experimental data and computed AP-scores for quantitative model. If other is not indicated, all metrics were averaged across features and cell types.

#### Simons Simplex Collection dataset processing and analysis of genomic variant effects

We have retrieved a set of variants discovered via whole genome sequencing of 1902 quartet families from the Simons Simplex Collection (supplementary table 2 in An et al. 2018). Only non-indel proband variants in the 50Kbp proximity of protein-coding genes’ transcription starts (as per GENCODEv38) were used. The dataset was lifted over from GRCh38 to GRCh37 human reference genome, resulting in 50 155 unique variants. We also retrieved another set of variants found independently in 1790 families from the same collection (supplementary table 1 in Zhou et al. 2019). These were processed by the same rules except for lifting over (as coordinates there were already on GRCh37), resulting in 28 869 unique variants.

Variants were used to modify sequence-tail input of DeepCT. We used DeepCT MPI model to infer signals of 40 epigenetic features in 794 cell types for each reference and alternative allele. We then subtracted predictions obtained for alternative allele from predictions obtained for reference allele and defined the obtained difference as cell type-specific *effect* of the variant on epigenetic feature. We next found maximum absolute effect across all cell types and features for each genomic variant. We’ve selected 500 variants with the biggest maximum absolute effect as a representation of a top 1% of the original variant set, found closest transcription start sites of protein-coding genes and submitted those to the PANTHER gene overrepresentation test (Fisher’s exact test with FDR correction, using Gene Ontology biological process annotation; (Mi et al. 2021, 16)).

Analysis of transcription factors motifs was performed using HOCOMOCO online tools (Kulakovskiy et al. 2018).

## Code availability

DeepCT code available at https://github.com/AIRI-Institute/DeepCT. Please note that DeepCT requires a custom version of selene framework, available at https://github.com/AIRI-Institute/selene

## Authors contribution

V.F., M.A., O.K., M.S. and A.L. conceived the study; M.A. implemented DeepCT with help from V.F., A.L., N.B., M.A. and E.M; N.C. and V.F. obtained and processed epigenetic datasets. N.C. and M.S. performed analysis of Simons Simplex Collection. O.K., V.F. and M.A. supervised the study. All authors contributed to manuscript preparation, read and accepted the final manuscript version.

## Acknowledgements

This study was performed using the infrastructure of AIRI, Artificial Intelligence Research Institute (Moscow, Russia). Preliminary analysis of ENCODE datasets was performed by Veniamin Fishman in the ICG (project No. 121031800061-7)

## Supplementary information

### Supplementary Note 1. Model architectures

#### A. Sequence tail of CNN models, layer parameters provided in bracket

Conv1d(4, 320, kernel_size=8)

ReLU

Conv1d(320, 320, kernel_size=8)

ReLU

MaxPool1d(kernel_size=4, stride=4)

BatchNorm1d(320)

Conv1d(320, 480, kernel_size=8)

ReLU

Conv1d(480, 480, kernel_size=8)

ReLU

MaxPool1d(kernel_size=4, stride=4)

BatchNorm1d(480)

Dropout(p=0.3)

Conv1d(480, 960, kernel_size=8)

ReLU

Conv1d(960, 960, kernel_size=8)

ReLU

BatchNorm1d(960)

Dropout(p=0.3)

Linear (42240, 256)

#### B. Cell type tail of the model

Linear (N_cell_types, 32)

#### C. Model’s head (same architecture used for regressor and classified). We used embedding_length equal 288 (concatenated sequence and cell type embeddings) or 256 (sequence embedding)

Linear(embedding_length, embedding_length)

ReLU

BatchNorm1d(embedding_length)

Linear(embedding_length, embedding_length)

ReLU

Linear(embedding_length, embedding_length)

ReLU

Linear(embedding_length, embedding_length)

ReLU

BatchNorm1d(embedding_length)

Linear(embedding_length, embedding_length)

ReLU

Linear(embedding_length, embedding_length)

ReLU

Linear(embedding_length, embedding_length)

ReLU

BatchNorm1d(embedding_length)

Linear(embedding_length, n_genomic_features)

**Supplementary Table 1.**
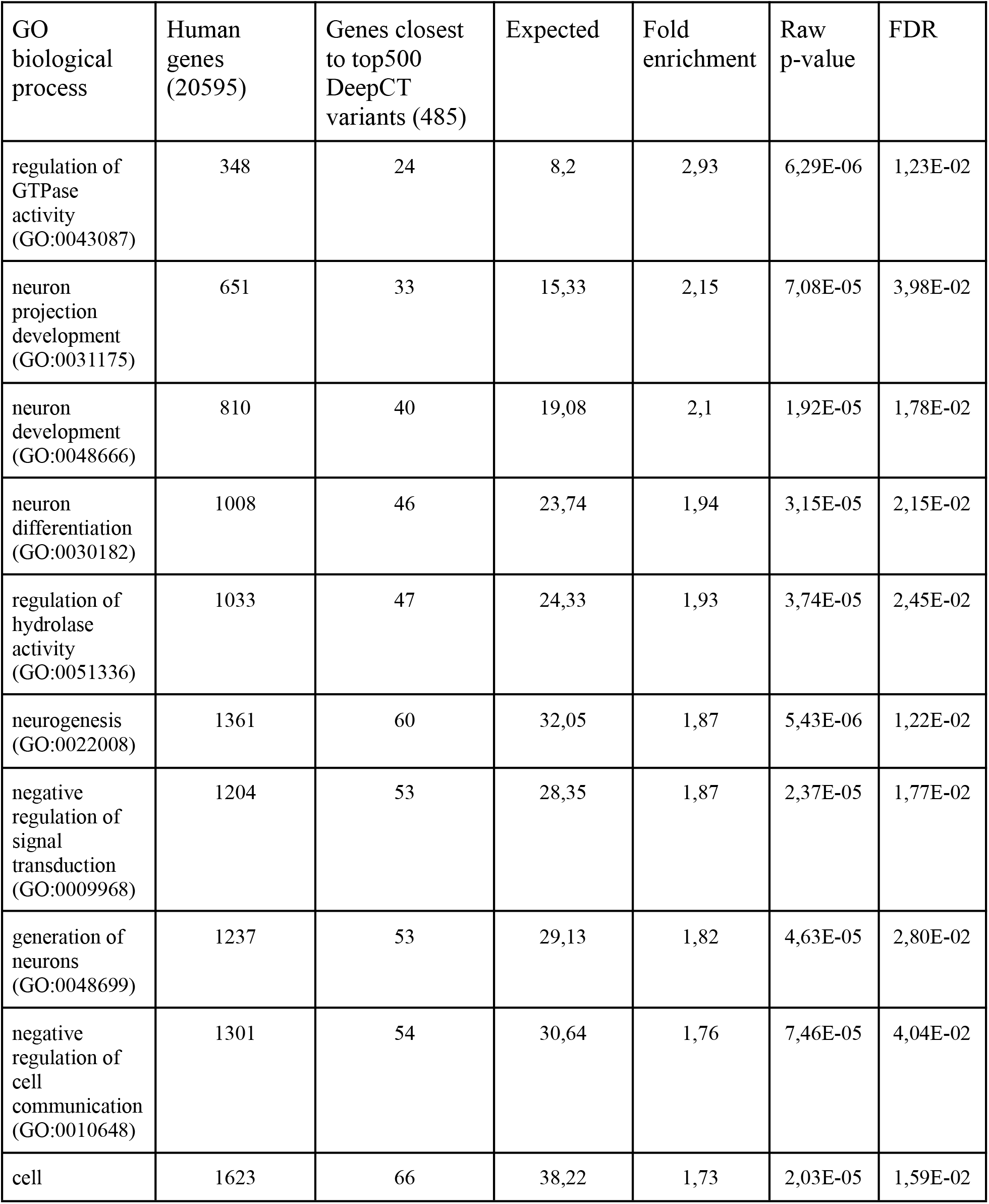
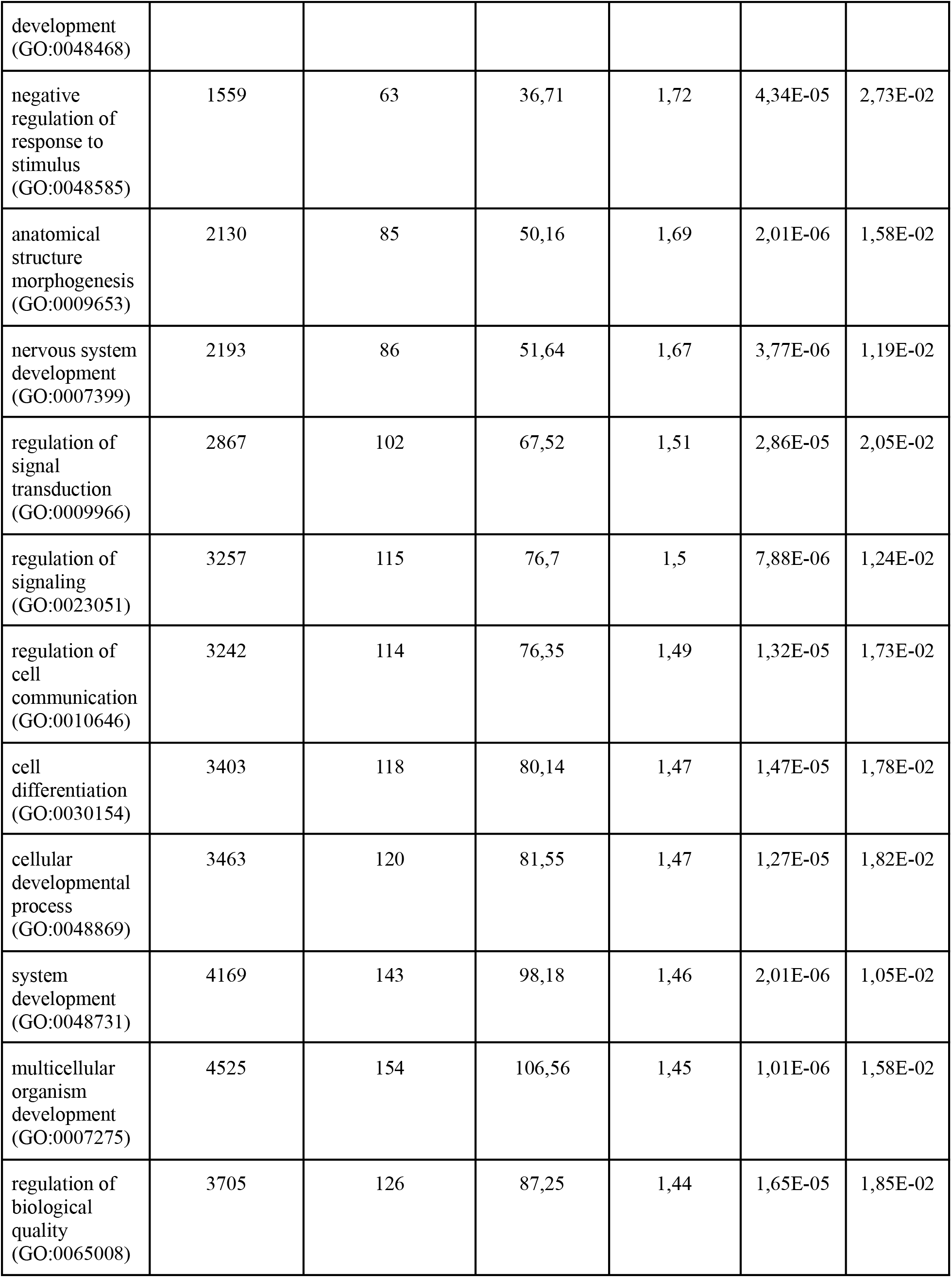
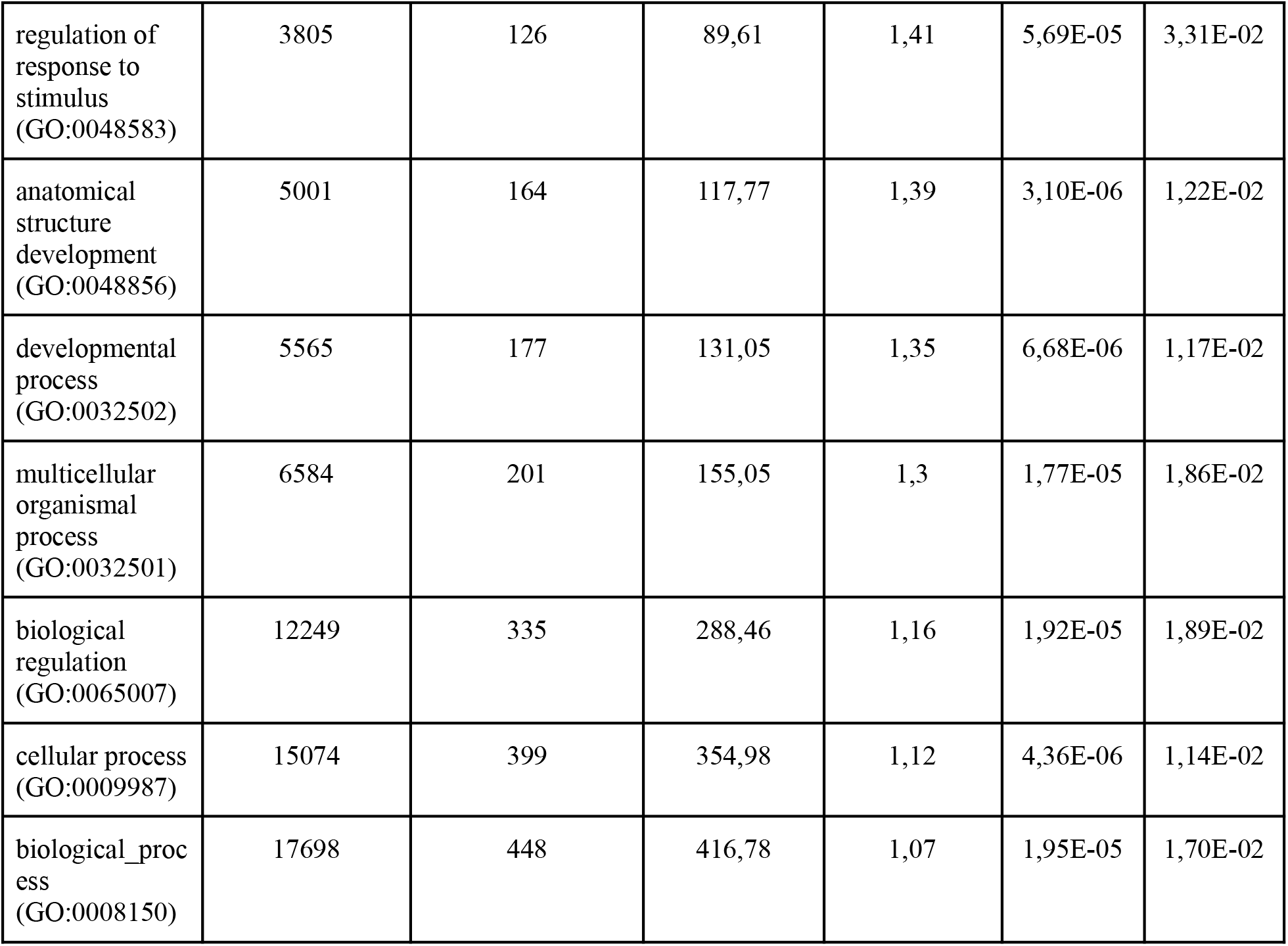
Results of gene enrichment analysis over the genes closest to 500 variants with the highest epigenetic effect sizes.

**Supplementary Figure 1.**
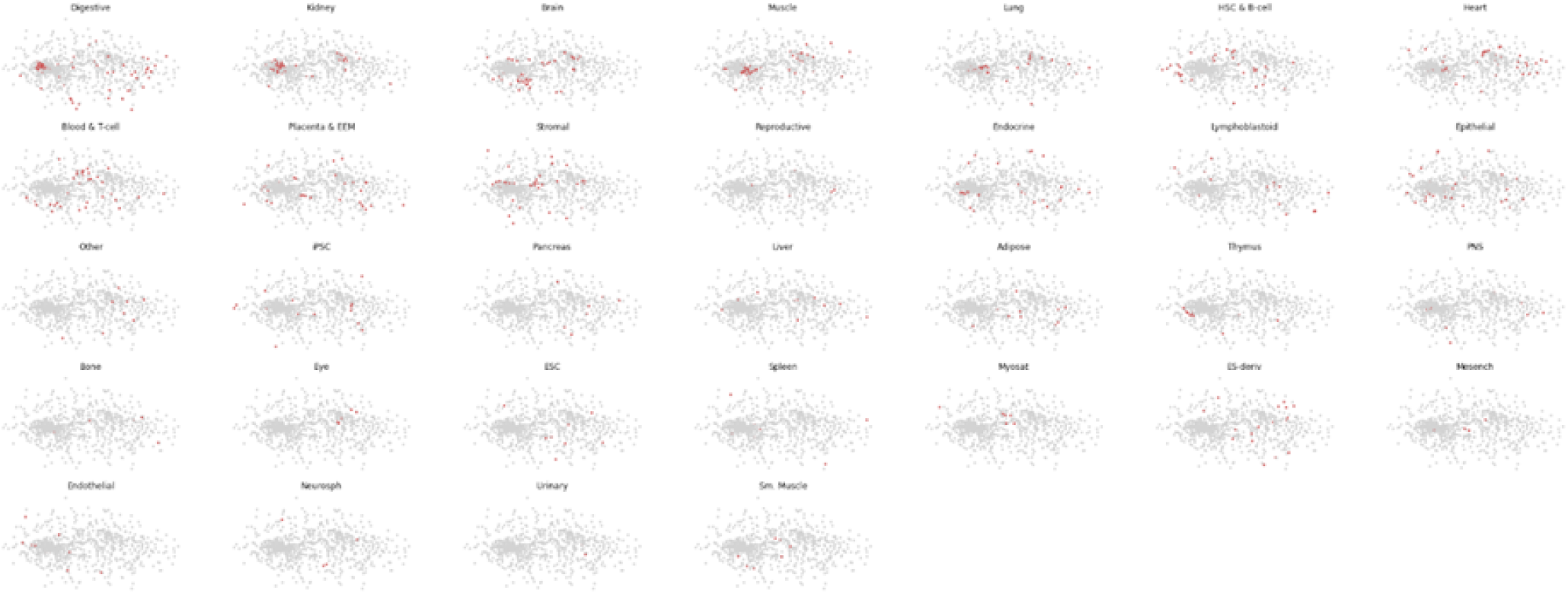
Cell state embeddings projections from 32-dimensional latent space of the model into 2-dimension PCA axis. Background points show distribution of embeddings for all cell types, whereas colored dots correspond to the specific cell types according to the subplot title.

**Supplementary Figure 2.**
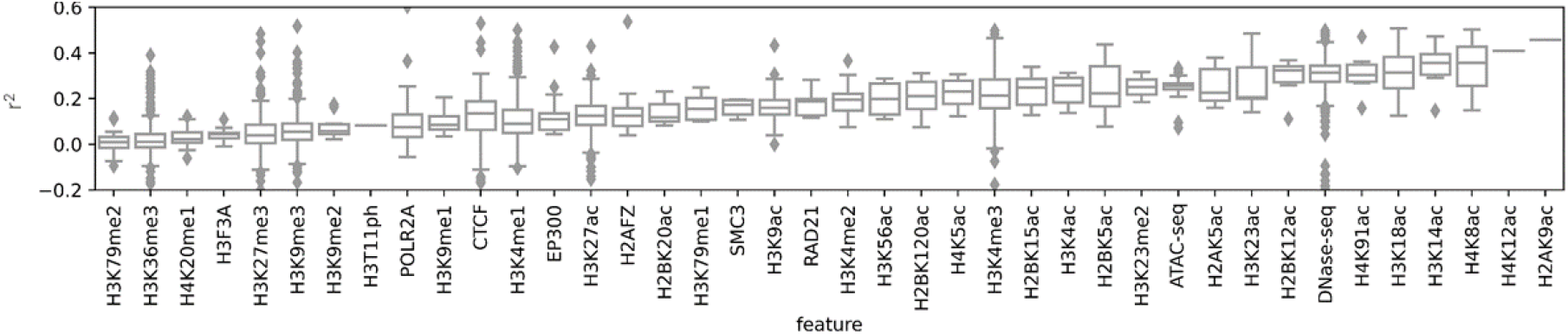
DeepCT performance in unseen track benchmark. Same as Fig. 3, D, but using r2 metrics.

**Supplementary Video 1**. *Supplementary video showing how model fits embeddings for three kidney and three muscle cell types. Each frame shows projection of embeddings from 32-dimensional latent space of the model into 2-dimension PCA axis*.

